# Nontypeable *Haemophilus influenzae* redox recycling of protein thiols promotes resistance to oxidative killing and bacterial survival in biofilms in a smoke related infection model

**DOI:** 10.1101/2021.08.25.457736

**Authors:** Benjamin C. Hunt, Xin Xu, Amit Gaggar, W. Edward Swords

## Abstract

Smoke exposure is a risk factor for community acquired pneumonia, which is typically caused by host adapted opportunists like nontypeable *Haemophilus influenzae* (NTHi). Genomic analyses of NTHi revealed homologs of enzymes involved in thiol metabolism, which can have key roles in oxidant resistance. Using a clinical NTHi isolate (NTHi 7P49H1), we generated isogenic mutant bacterial strains in which homologs of glutathione reductase (NTHI 0251), thiol peroxidase (NTHI 0361), thiol peroxidase (NTHI 0907), thioredoxin reductase (NTHI 1327) and glutaredoxin/peroxiredoxin (NTHI 0705) were inactivated. Bacterial protein analyses revealed significant increases in protein oxidation after oxidative stress for all the mutant strains. Similarly, each of these mutants were less resistant to oxidative killing compared with the parental strain; these phenotypes were reversed by genetic complementation. Quantitative confocal analysis of biofilms showed reducted biofilm thickness and density, and significant sensitization of bacteria within the biofilm structure to oxidative killing for thiol mutant strains. Smoke-exposed mice infected with NTHi 7P49H1 showed significantly increased lung bacterial load, as compared to control mice. Immunofluorescent staining of lung tissues showed NTHi communities on the lung mucosa, interspersed with host neutrophil extracellular traps; these bacteria had surface moieties associated with the *Hi* biofilm matrix, and transcript profiles consistent with NTHi biofilms. In contrast, infection with the panel of NTHi mutants showed significant decrease in lung bacterial load. Comparable results were observed in bactericidal assays with neutrophil extracellular traps in vitro. Thus, we conclude that thiol mediated redox homeostasis promotes persistence of NTHi within biofilm communities.

**Importance:** Chronic bacterial respiratory infections are a significant problem for smoke exposed individuals, especially those with chronic obstructive pulmonary disease (COPD). These infections often persist despite antibiotic use. Thus, the bacteria remain and contribute to the development of inflammation and other respiratory problems. Respiratory bacteria often form biofilms within the lungs, while growing in a biofilm their antibiotic and oxidative stress resistance is incredibly heightened. It is well documented that redox homeostasis genes are upregulated during this phase of growth. Many common respiratory pathogens such as NTHi and *Streptococcus pneumoniae* are reliant on scavenging from the host the necessary components they need to maintain these redox systems. This work here begins to lay down the foundation for exploiting this requirement and thiol redox homeostasis pathways of these bacteria as a therapeutic target for managing chronic respiratory bacterial infections, which are resistant to traditional antibiotic treatments alone.

## Introduction

Cigarette smoke exposure, be it primary or secondary, extracts a significant economic and health burden in the United States and globally. In the US alone, a significant proportion of healthcare expenditures are directed at dealing with smoke related issues (1). Cigarette smoke is a complex mixture that elicits significant chances in vascular, airway, and immune function, which lead to the development and exacerbation of a variety of diseases (2–8). Exposure to cigarette smoke is a significant risk factor for community acquired pneumonia with both bacterial and viral infections that can exacerbate and further contribute to the development of smoke associated morbidities (2, 9–11).

Smoking is associated with development of chronic bacterial infections which are typically caused by host adapted opportunists such as nontypeable *Haemophilus influenzae* (NTHi) (12–16). NTHi is a gram-negative pathobiont that typically asymptomatically resides in the nasopharynx with little to no overt pathology. However, in persons with compromised airway clearance, NTHi can colonize the lower airways and establish chronic infections (12, 16–18). NTHi are thought to persist within biofilm communities on the airway mucosa during a variety of opportunistic infections (19). Biofilms are complex, heterogenous communities that are intransient to environmental stressors, antibiotics, or host immune effectors largely due to so-called “persister” subpopulations within the biofilm structure (20–22). These bacterial biofilms display unique genetic expression profiles, enhanced antimicrobial and oxidative stress resistance, and increased resistance to immune cell clearance compared to planktonically growing bacteria (18, 20, 23–26).

Phagocytes undergo an “oxidative burst” that culminates in release of reactive oxygen species (ROS), which is a key component of the innate immune response to bacterial infection (27, 28). Thus, maintaining proper redox homeostasis and having mechanisms for counteracting oxidative stress is vital for pathogens to colonize and persist within their host (29–32). Bacteria respond to ROS by activating antioxidant defenses, shifting metabolic pathways, and promoting the formation of biofilms (30, 33, 34). Glutathione (GSH) is a cysteine containing thiol tripeptide with important roles in oxidative stress defenses in a wide array of biological systems (29, 35–37). Importantly, *Streptococcus pneumoniae* and *Haemophilus influenzae* are reliant on the import of exogenous GSH from the airway environment where it is abundant (38). Peroxiredoxin/glutaredoxin (*pdgX*) is a GSH metabolic enzyme which has been shown to be upregulated within NTHi biofilms, as well as patient sputa (15, 29, 30).

Analysis of sequenced NTHi genomes revealed a number of homologs of enzymes involved in reduction of oxidized thiols, including predicted glutathione reductase (*gor*, NTHI0251), thiol peroxidase (*bcp*, NTHI0361), thiol peroxidase (*tpx*, NTHI0907), thioredoxin reductase (*trxB*, NTHI1327) and glutaredoxin/peroxiredoxin (*pdgX*, NTHI0705) (**Table 1**). We used a bacterial genetic approach to generate a number of isogenic mutant strains which were then used investigate the importance of this pathway in colonization, persistence, and biofilm formation within the airways. We show that disruption of thiol redox homeostasis in NTHi results in significant susceptibility to oxidative stress, neutrophil extracellular trap killing, and defects in persistence in a susceptible smoke exposed mouse infection model. Based on these results, we conclude that thiol metabolism is an important determinant of NTHi colonization and persistence within biofilm communities.

**Table 1.** Gene designations and predicted functions based on homology for genes of interest. Allele numbers are given in parentheses.

| Glutathione/ Biofilm Related Genes of Interest |  |
| --- | --- |
| Gene | Predicted Gene Function |
| *gor* (NTHI0251) | Predicted glutathione-disulfide reductase |
| *grxA* (NTHI1601) | Glutaredoxin |
| *bcp* (NTHI0361) | Thioredoxin-dependent thiol peroxidase |
| *trxB* (NTHI1327) | Thioredoxin-disulfide reductase |
| *pgdX* (NTHI0705) | Peroxiredoxin-glutaredoxin |
| *tpx* (NTHI0907) | Thiol peroxidase |
| *luxS* (NTHI0621) | AI-2 synthesis protein, quorum signaling |
| *dps* (NTHI1817) | DNA binding ferritin-like protein |

## Materials and methods

### Bacteria and culture methods

NTHi 7P49H1 (provided by Dr. Timothy Murphy, University of Buffalo) is a sputum isolate from a patient with chronic obstructive pulmonary disease (39). NTHi bacteria were cultured at 37° C on brain heart infusion agar (Difco, NJ, USA) supplemented with nicotinamide adenine dinucleotide (10 μg/ml, Sigma) and hemin (10 μg/ml, ICN Biochemical). Bacteria were harvested from the surface of overnight culture plates and resuspended in PBS to the desired optical density to generate inocula.

### Measurement of bacterial resistance to oxidant

Bacteria were suspended in sBHI media and seeded at a concentration of ~10^8^ CFU/ml into a 24-well dish, cultured at 37° C and 5% CO_2_ for 24 h, after which growth media was aspirated and replaced with PBS containing varying concentrations of hydrogen peroxide as indicated in figure legends. Bacteria were exposed to oxidant for 30 min after which surface adherent bacteria were gently washed with PBS three times. Bacterial biofilms were then scraped off the bottom of the well, serially diluted, and plated on sBHI for plate-count.

### Quantification of cysteine sulfenic acid oxidation

Bacteria were resuspended in PBS to an ~10^8^ CFU/mL; bacterial density was confirmed by plate-count. Bacteria were then centrifuged, and the supernatant was removed and replaced with a 500 mM hydrogen peroxide solution, and incubated for 30 minutes at 37°C, after which the supernatant was removed and pellet washed with PBS. Bacteria were lysed enzymatically using a lysis buffer (50 mM Tris pH 8.0, 10% glycerol, 0.1% TritonX-100, and 100 mg/ml lysozyme), while simultaneously labeling for cysteine sulfenic acids (CSAs) with 1mM biotin-1,3-cyclopentanedione (BP1) (Kerafast, Boston, MA, USA). Lysis buffer was prepared fresh before use and BP1 was added immediately prior to use. Samples were lysed and labeled for CSAs for 1 hour at 37°C. After lysis and biotin labeling of CSAs, the CSA modified proteins were isolated using Takara Bio Capturem^™^ streptavidin miniprep columns (Shiga, Japan) following the manufacturer’s instructions. Next, samples were run on an 4-20% gradient SDS-PAGE gel, stained using the Pierce^™^ silver stain kit, and imaged. Relative CSA protein modifications was determined via densitometry using ImageJ Fiji (40).

### Measurement of bacterial killing by neutrophil extracellular traps

Derivation and measurement of NTHi killing by neutrophil extracellular traps was essentially as described in previous studies (25, 41). HL60 monocyte cells were cultured in RPMI 1640 (Thermofisher, MA, USA) with 10% FBS, and differentiated for 5-6 d in RPMI containing 0.8% dimethyl formamide, then collected by centrifugation. Cells (~10^6^/ well) were seeded into wells of a 24-well dish and activated with 25 nM phorbol myristate acetate (PMA) (Sigma-Aldrich, MO, USA) for 10 minutes. NET formation was confirmed via microscopy. Cell culture media, with or without 20 μM cytochalasin D, was then added and cells were incubated for 15 minutes. Bacterial strains were then added at an MOI of 10 in triplicate and incubated for 30 minutes, then scraped and serially diluted and plated to enumerate bacterial counts. Bacterial killing was expressed as percentage of counts obtained from control wells with no HL60 cells. NET vs phagocytic killing was obtained by comparison to wells with cytochalasin D to those without.

### Cigarette smoke mxposure and mouse infections

Mice (C57BL/6J) were acquired from Jackson Laboratory (Bar Harbor, ME, USA) and randomly assigned to an experimental group, whole cigarette smoke or ambient air control. Smoke group mice were placed in a whole-body exposure chamber and exposed to smoke from 3R4F research cigarettes (Louisville, KY, USA), twice daily for a period of 14 days. Between the two smoke exposures for each day, animals were allowed a 2-hour rest period. Animals began smoke exposure at the minimum of 6 cigarettes per day, with the total number of cigarettes increase by 2 per day until reaching 24 total per day then remaining constant for the remainder of the regime. Animals were monitored continuously during smoke exposure. Cigarette smoke was generated by an automated cigarette smoke generator (SCIREQ, InExpose model), with a 24-cigarette carousel. SCIREQ filters were monitored and weighed to measure total particulate matter exposure for comparisons to other murine smoke models. Animals exposed to smoke were stored separately from the air control groups. After completing the smoke exposure regimen, mice intratracheally infected with 10^7^ CFUs of NTHi or vehicle control (PBS). Animals were euthanized 24, 48, and 72 hours post infection. All animal and infection procedures were performed according to AVMA laboratory standard procedures and were reviewed and approved by the UAB Institutional Animal Care and Use Committee.

### Immunofluorescent staining and confocal laser scanning microscopy

Mouse lung tissue sections were sectioned via cryostat (Thermofisher CryoStar NX70 cryostat, 5 μm/section) and fixed onto glass slides. NTHi bacteria were stained using polyclonal rabbit antiserum and goat anti-rabbit IgG Alexa 488 secondary antibody conjugate. Cover slips were mounted using Prolong Gold antifade reagent with DAPI (Thermofisher, Waltham, MA). Confocal laser scanning microscopic analyses were performed using a Nikon A1R TE2000 inverted microscope (Nikon, Tokyo, Japan). CellROX Deep Red was utilized, following manufacturer’s instructions, to image and measure oxidative stress in fixed bacterial biofilm samples (Thermofisher, Waltham, MA). Pixel intensity maps and quantification of biofilm images was done using BiofilmQ software (42). Images were segmented using the semi-manual Otsu threshold method. NTHi sialylated moieties in biofilm matrix were stained utilizing specific lectin conjugates, essentially as described previously (43). *Maackia amurensis* lectin (MAA) Texas Red conjugated specific for Neu5Ac α(2,3) galactose (EY Laboratories, San Mateo, CA) was diluted to a final concentration of 100 μg/ml in 0.01 M phosphate, 0.15 M NaCl, 0.05 M sodium azide buffer according to manufacturer’s instructions. Representative images were created using Fiji imaging analysis software (40).

### Transcript quantification using qRT-PCR

Bacterial RNA was extracted using the Monarch total RNA miniprep kit (New England Biolabs, Ipswich, MA) following manufacture’s guidelines. RT-qPCR was performed using the Applied-Biosystems 7500 System, and oligonucleotide probes specific for *pdgX*, *luxS*, *dps*, *hktE* and *omp26* (**Table 2**). The *omp26* transcript was chosen as an endogenous control given that its expression does not vary between planktonic or biofilm mode of growth (44). The reaction mix used was the NEB Luna Universal following manufacturer’s directions for cycling conditions (New England Biolabs, Ipswich, MA). All samples were run in duplicate. Transcript measures were normalized relative to *omp26* levels from the same sample. Relative quantification of gene expression was determined using the comparative CT method (2^ΔΔCT^).

**Table 2.**
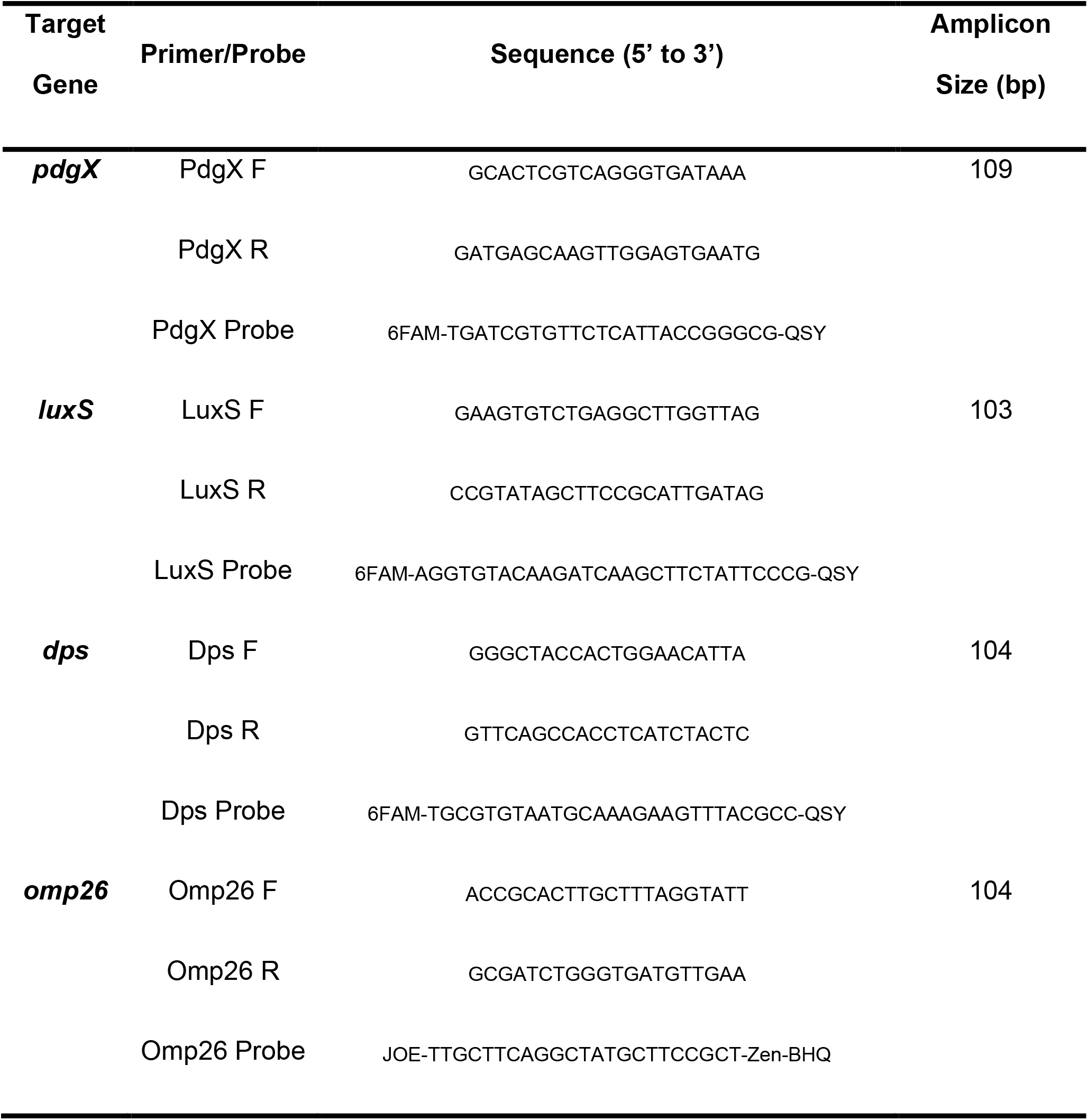
RT-qPCR primers and probes used for analysis of biofilm expression genes.

### Statistical analyses

Data were analyzed by the one-way analysis of variance (ANOVA) with Tukey’s multiple comparisons test. All bar graphs represent mean and error bars represent standard error of the mean (SEM) or standard deviation (SD) as noted in the figure legend. All mouse infection experiments were repeated at least three times. Data from multiple independent animal experiments were averaged together. Animal numbers in each group are denoted in the figure legends. * P < 0.05, ** P <0.005, *** P <0.0005, **** P <0.00005; *P* values < 0.05 were considered statistically significant.

## Results

To assess the susceptibility of NTHi 7P49H1 and our isogenic mutant strains to oxidative stress, we performed a killing assay where we exposed NTHi biofilms to varying concentrations of hydrogen peroxide (H_2_O_2_) for 30 min. Similar to prior experiments with other NTHi strains, minimal killing of NTHi 7P49H1 was observed except at the highest concentrations of hydrogen peroxide. In contrast, each of the isogenic mutant strains had significant levels of killing with bacterial counts going below the detectable limit of viable plate counting when exposed to hydrogen peroxide. Resistance to oxidant was restored by genetic complementation of the isogenic mutants (Figure 1A). Comparable results were obtained in parallel experiments with hypochlorous acid (HOCl) as the oxidative stressor (data not shown).

**Figure 1.**
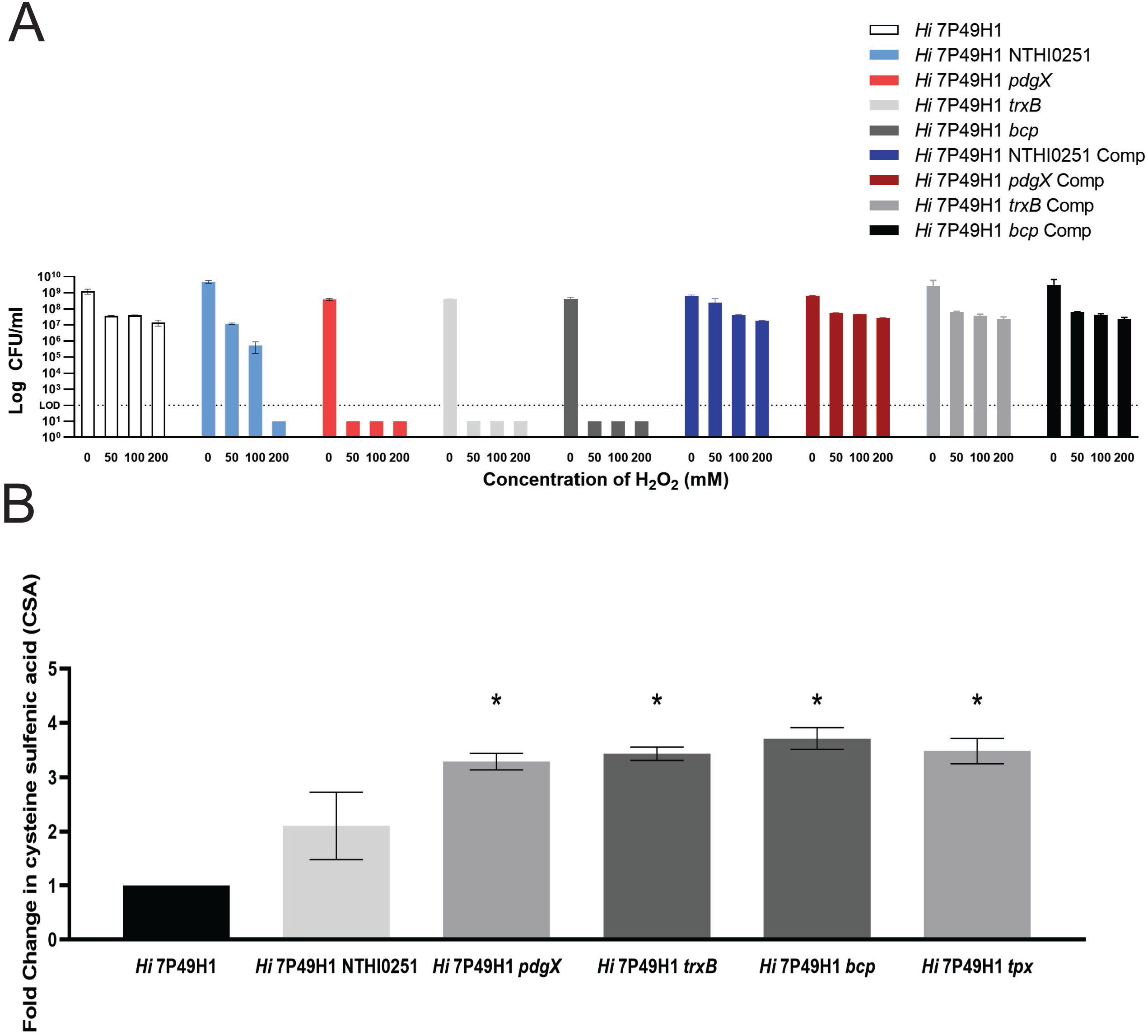
Bacterial susceptibility to varying concentrations of hydrogen peroxide (H_2_O_2_) and quantification of cysteine sulfenic acid protein modifications. A) Oxidative stress and killing of parent, thiol mutant, and complemented strains as determined by viable plate counting post exposure. The dashed line represents the limit of detection (LOD). Mean +/- SEM (N=5). Data is representative of three biological replicates. B) Fold change in cysteine sulfenic acid (CSA) protein modifications post exposure to 500 mM hydrogen peroxide as determined by densitometry. Mean +/- SD (N=3).

To further investigate the susceptibility and redox state of the isogenic mutants, we purified oxidized bacterial proteins based on affinity of cysteine sulfenic acid (CSA) for the nucleophile 1,3-cyclopentanedione (BP1) (45–47). Biotin-linked BP1 was used to purify CSA-modified proteins from bacterial lysates, which were quantified by SDS/PAGE and silver stain. In comparison to the parent strain, thiol redox mutants displayed significantly higher levels of cysteine sulfenic acids and displayed 2-to-3.5-fold higher change in cysteine sulfenic acid levels (Figure 1B). This is indicative of an imbalance in the redox homeostasis of the isogenic mutants and of a heightened sensitivity to oxidative stress. Together, reactive oxygen species killing and the measurement of cysteine sulfenic acids show that disruption of the thiol redox pathway at different steps can similarly sensitize and compromise NTHi’s ability to maintain redox homeostasis.

To further investigate the redox homeostasis and oxidant resistance of our isogenic mutant biofilms when exposed to oxidative stress, we utilized confocal scanning laser microscopy alongside a fluorescent stain to visualize the areas of the biofilm with the highest degree of reactive oxygen species and BiofilmQ to quantitatively analyze the confocal images. Using confocal imaging, we generated vertical Z-series NTHi biofilms after exposure to 500 mM hydrogen peroxide, with the objective of defining any effects on biofilm formation/maturation and identify and localize bacterial subpopulations differentially impacted by oxidative stress. As observed previously, NTHi formed thick communities extensive three dimensional height and structure, which are maintained after exposure to oxidative stress. In contrast, NTHi 7P49H1 *pdgX* biofilms were largely diminished in three-dimensional structure following peroxide treatment, consistent with reduced resistance (Figure 2A).

**Figure 2.**
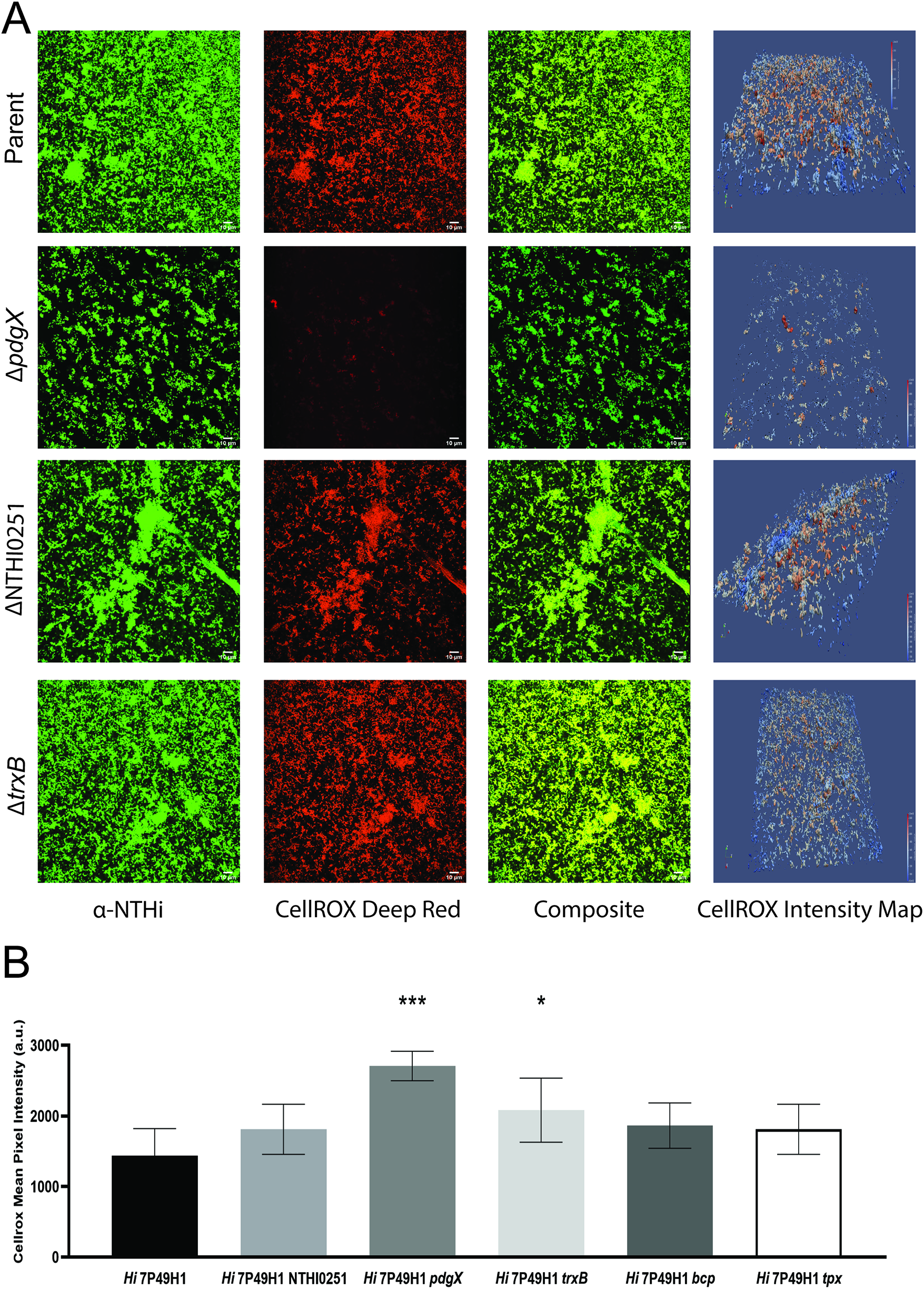
Immunofluorescent imaging and analysis of redox stress of *in vitro* NTHi biofilms. A) Immunofluorescent staining of 24 hour biofilms exposed to 500 mM hydrogen peroxide. Bacteria were stained with anti-NTHi antibodies conjugated to Alexa 488 (green) while bacterial oxidative stress was visualized using CellROX Deep Red (red). Fluorescent pixel intensity maps of the CellROX channel were generated using BiofilmQ. Images were taken at 60X magnification. Scale bars represent 10 μm. B) Mean pixel intensity of the CellROX channel was quantified using BiofilmQ software. Mean +/- SD (N=5).

Quantifying and localizing areas with highest intensity of CellRox fluorescent signal, and thus presumably higher oxidative stress, were the areas of the biofilm closest to the substrata and the internal areas of mature tower structures (Figure 2A). Additionally, when utilizing the BiofilmQ software to quantitate the mean pixel intensity of the CellRox fluorescent signal, it was found that the isogenic mutants displayed heightened mean pixel intensities. Of the isogenic mutant strains, NTHi 7P49H1 *pdgX* and NTHi 7P49H1 *1327*, were found to have significantly higher mean pixel intensities when compared to the parent strain. The NTHi 7P49H1 *pdgX* isogenic mutant displayed the highest mean intensity, nearly double the mean for the parent strain (Figure 2B). This indicates that the isogenic mutant biofilms are inherently more susceptible to oxidative stress than the parent strain, particularly the NTHi 7P49H1 *pdgX* isogenic mutant biofilm.

Our *in vitro* data spurred on the desire to establish and utilize a smoke exposed murine model and investigate the consequences of the disruption of redox homeostasis on persistence and disease. To assess the impact of thiol metabolism on NTHi colonization and persistence in the lung, we performed infection studies on mice (C57/Bl6) following exposure to cigarette smoke. The smoke exposure regime and timeline of infections are outlined in Figure 3A. Bacteria were recovered from the lungs of smoke exposed mice infected by NTHi 7P49H1 for up to 48 h post-infection; this was in contrast to control mice which cleared infection. Each of the thiol redox mutants tested were unable to establish a successful infection in the susceptible smoke exposed mouse lung, showing no detectable amounts of NTHi at any time post infection (Figure 3B). Animals infected with isogenic thiol mutants had significantly less weight loss over the course of infection, indicating overall less severe disease.

**Figure 3.**
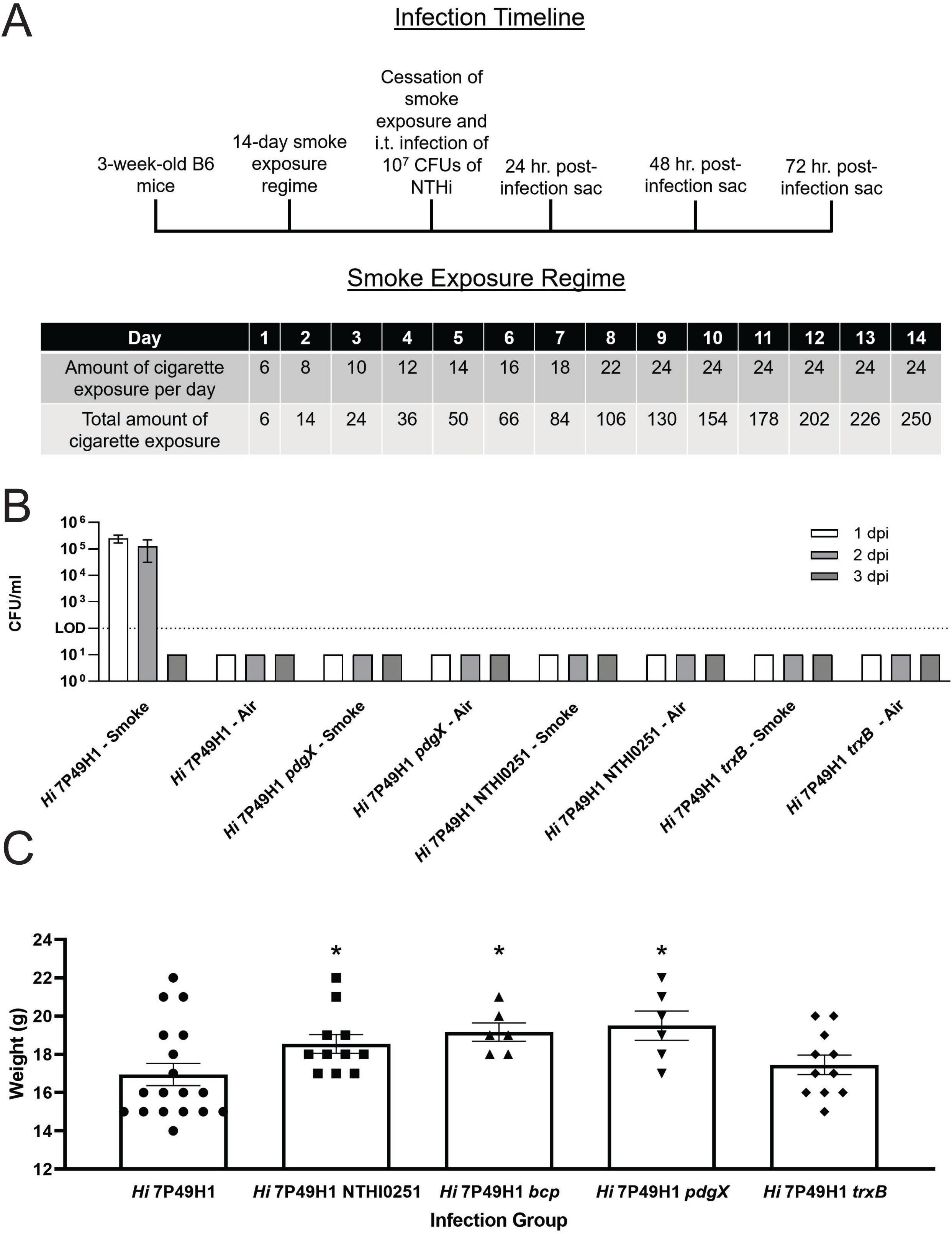
Bacterial infection of smoke exposed and air control murine lungs with NTHi and thiol redox mutants. A) Schematic detailing infection timeline and smoke exposure regimen for mice. B) Persistence in smoke exposed and air control murine lungs as determined by viable plate counting. Dotted line indicates limit of detection (LOD). Mean +/- SEM (N=6-18). C) Weight of mice 24 hours post infection. All comparisons of weight were made to the parent strain infected animal group. Mean +/- SEM (N=6-18). Data is representative of three independent biological replicates.

Using confocal scanning laser microscopy revealed that in the airways of smoke exposed mice infected with the parent strain multicellular NTHi communities can be detected at 24 and 48 hours post infection (Figure 4A). These some multicellular communities are absent in the airways of animals infected with isogenic thiol mutants. To confirm that these multicellular communities found within the airways were NTHi biofilms stained the tissue sections with fluor conjugated lectins that would bind specific linkages within the NTHi biofilm extracellular matrix. Doing so revealed that our aggregates of NTHi overlap with the staining for the biofilm extracellular matrix, thus showing that these communities are indeed imbedded in a biofilm like structure (Figure 4A). Additionally, RT-qPCR analysis of bacterial transcripts isolated from the airways 48 hours post infection revealed that the expression of biofilm related genes (*pdgX*, *luxS*, *dps*) were elevated. Combined with the confocal imaging, the expression of biofilm associated genes further cements that these structures are indeed NTHi biofilms forming within the lungs of susceptible smoke exposed mice.

**Figure 4.**
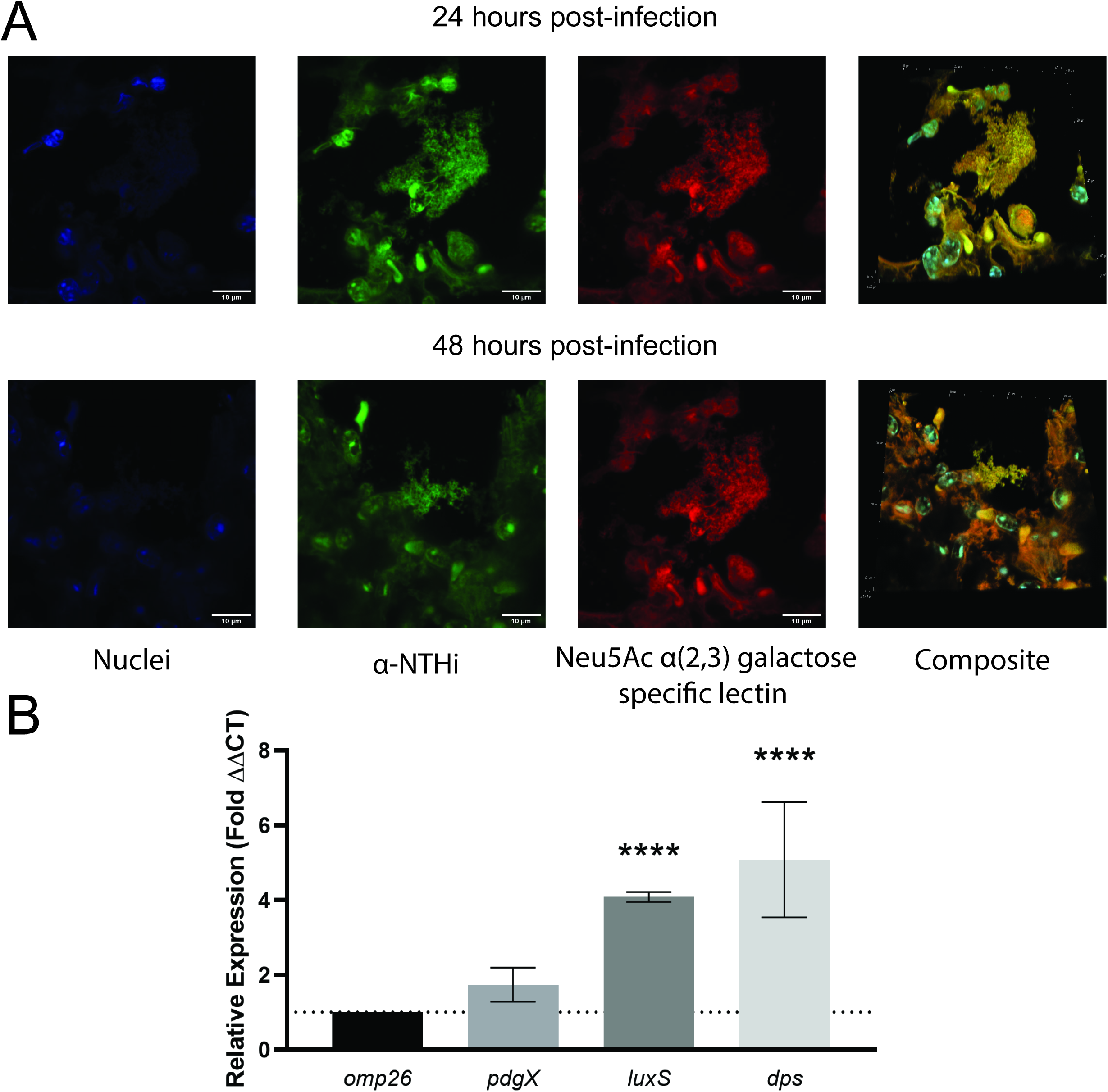
Immunofluorescent imaging and RT-qPCR analysis of NTHi biofilms in the smoke exposed murine lung. A) Immunofluorescent staining of parent strain infected mouse lung sections. Lung sections are representative of 24 and 48 hours post infection. Nuclei were stained with DAPI (blue) and bacteria were stained with anti-NTHi polyclonal antibodies conjugated to Alexa 488 (green). NTHi biofilm components were stained using the *Maackia amurensis* lectin (MAA) conjugated to Texas-Red, which is specific for neu5Ac α(2,3) galactose linkages (red). Images are taken at 90X magnification. Scale bars represent 10 μm. B) RT-qPCR analysis of mRNA isolated from infected mouse lung of biofilm associated genes, in comparison to *omp26* housekeeping gene. Samples were collected from 24 hours post infection. All samples were run in triplicate. Mean +/- SD N=4-6.

Prior work from our group has shown that resistance to oxidant is central NTHi survival and growth within neutrophil extracellular traps and thus important for establishing a chronic infection (25). To assess bacterial survival of our thiol redox isogenic mutants and to gain deeper insight into the lack of persistence within our smoke exposed murine model we turned to cell culture methods and confocal scanning laser microscopy methods once again. To investigate the susceptibility of our mutant strains to killing via neutrophil extracellular traps we used differentiated Hl60 cells activated with phorbol myristate acetate and inhibited phagocytosis using cytochalasin-D. Unsurprisingly, the parent strain was highly resistant to killing via neutrophil extracellular traps, whilst the isogenic thiol mutant strains were each significantly more susceptible to killing (Figure 5A). Curious about this finding that our mutant strains had a heightened susceptibility to neutrophil extracellular trap killing; we returned to our lung tissue sections and confocal scanning laser microscopy, however, this time we stained for DNA and citrullinated histone H3, both components of neutrophil extracellular traps. We see in the airways of smoke exposed mice 48 hours post infection NTHi multicellular communities surrounded by neutrophil extracellular traps (Figure 5B). Additionally, using a myeloperoxidase activity assay to show the increased level of myeloperoxidase activity at 48 hours post infection combined with the presence of neutrophil extracellular traps, these surrounding neutrophils are activated (Figure 5C). Together, these data showing the susceptibility of our isogenic thiol strains to killing via neutrophil extracellular traps and the finding that NTHi biofilms in our smoke exposed murine airways are surrounded by neutrophil extracellular traps indicates that the isogenic thiol mutant strains might have failed to establish a persistent infection due to these deficiencies.

**Figure 5.**
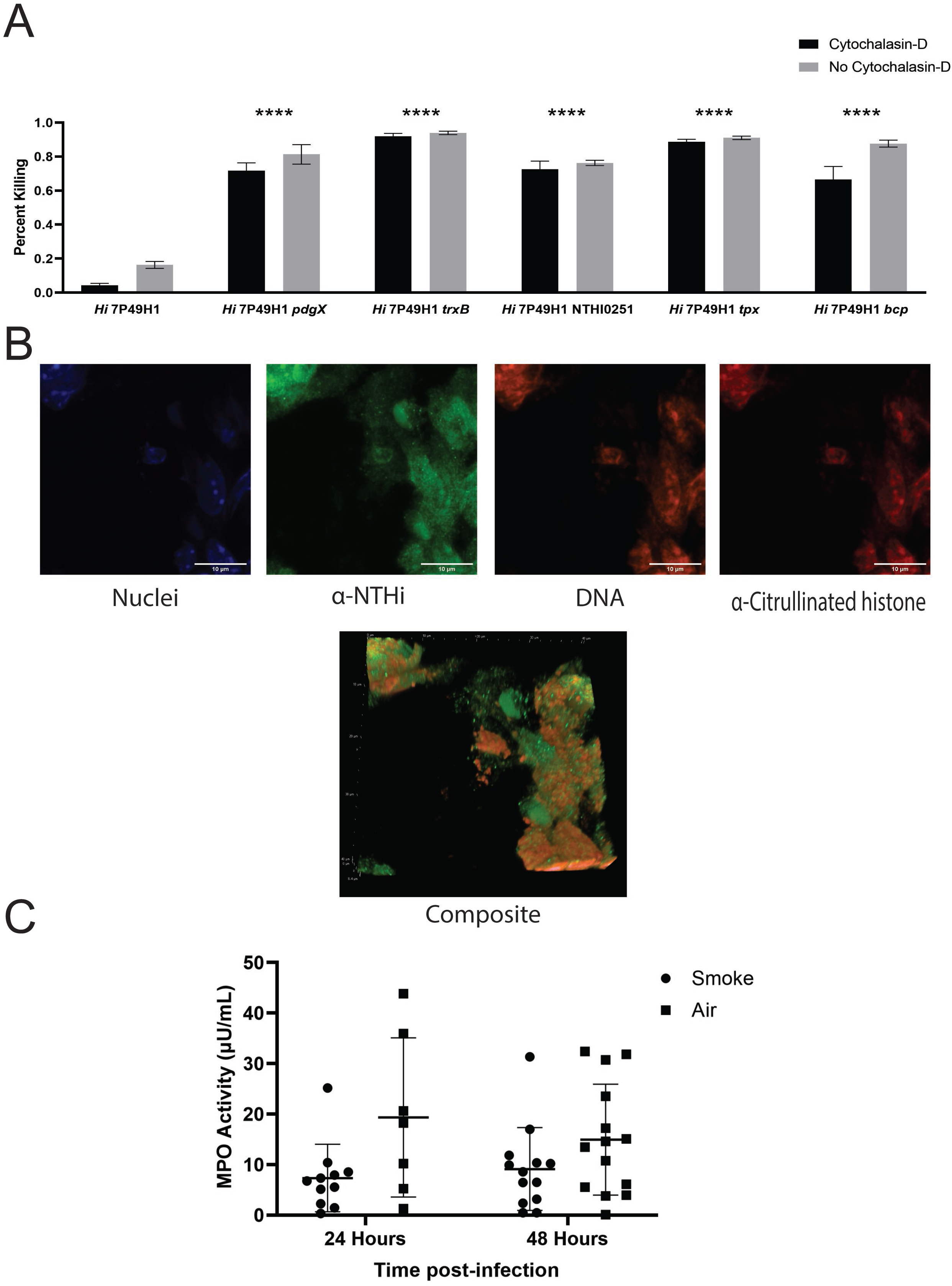
Neutrophil extracellular trap (NET) killing assay and fluorescent staining for NETs in infected mouse lungs. A) 5-6 day differentiated HL60 cells seeded in 24-well plates were activated with 25 nM phorbol myristate acetate after which 20 μM cytochalasin-D was added to prevent killing via phagocytosis. Bacteria were added at an MOI of 10. Bacterial killing was expressed as a percentage of counts obtained from control wells. Mean +/- SEM (N=3). All statistical comparisons are made to the matched treatment of the parent strain. B) Immunofluorescent staining and confocal imaging of infected mouse lung sections. Nuclei were stained with DAPI (blue), bacteria were stained with anti-NTHi polyclonal antibodies conjugated to Alexa 488 (green), while NETs were stained using anti-Histone H3 Citrulline R2, R8, R17 (red) and propidium iodide (orange). These confocal images are representative images of mouse lungs 48 hour post infection. Images are taken at 90X magnification. Scale bars represent 10 μm. C) Myeloperoxidase (MPO) activity within the lungs of infected mice during the course of infection Mean +/- SD (N=12-18).

## Discussion

Chronic bacterial infections are a significant problem for patients with mucosal clearance defects, including patients with cystic fibrosis, COPD and preceding viral infection (16, 48, 49). Bacterial biofilms, which display heightened resistance to antibiotics, oxidative stress, and house persister cells, contribute significantly to the development of chronic infections (24, 30, 49–51). Thus, management of biofilm related infections and developing novel therapeutics to counteract the persistence advantages biofilms provide has the potential to greatly benefit many patient populations. It has been documented that a biofilm mode of growth, both *in vitro* and *in vivo*, is associated with an increase in the expression of redox related genes, one example being *pdgX*, which has been shown to be upregulated in the COPD lung (15, 30, 33, 44). Compromising a biofilm’s ability to respond to oxidative stress should render the bacteria within more susceptible to clearance by the ROS naturally generated by the immune system.

Before the development of novel therapeutics can begin, work must be done to investigate the mechanisms of thiol redox homeostasis that is employed by bacteria, in this instance we focus on the relevant pathogen nontypeable *Haemophilus influenzae*. Using a series of congenic mutants, we aim to investigate the impact of the disruption of thiol redox homeostasis at varying points in the pathway and the resulting consequences of doing so. Using *in vitro* analysis of the redox homeostasis capabilities of our mutants, we show that disruption along this pathway results in significant susceptibility to oxidative stress and excess protein damage (Figure 1A). Interestingly, disruption of these differing thiol redox genes resulted in similar phenotypes. This highlights the importance of this pathway and signifies that there are potentially multiple therapeutic targets within this pathway that are worth pinpointing. Additionally, using confocal scanning laser microscopy alongside BiofilmQ analysis, we were able to show that the mature biofilms of our mutant strains are susceptible to oxidative stress, with much of the highest levels of stress being located in the portions of the biofilm closest to the substrata (Figure 2).

Seeking to further delineate the *in vitro* phenotype, we utilized cell culture methods to quantify defects in immune cell resistance, specifically to neutrophil extracellular traps. Given the heightened susceptibility to oxidative stress our thiol redox mutants possess, it is unsurprising that our mutant strains are also significantly more susceptible to killing via neutrophil extracellular traps released from activated HL60 cells (Figure 5A). This data is encouraging and lends credence to the notion that if a therapeutic drug were able to disrupt NTHi thiol redox homeostasis, it could enable clearance by the normal immune response to NTHi.

The results discussed thus far have all been *in vitro*, however, using a smoke exposed murine model, we were able to investigate whether the same phenotypes persist when brought into a disease relevant *in vivo* model for NTHi lung infections (Figure 3A). As is the case in human patients, smoke exposed animals were susceptible to infection by the parent strain, however, thiol redox mutant strains were unable to establish an infection in the susceptible smoke exposed airways, showing no detectable CFUs at 24- or 48-hours post-infection (Figure 3B). Based on the *in vitro* data, it is reasonable to surmise that this may be due to defects in the ability to respond to the ROS produced by the immune system during infection. Additionally, using confocal microscopy and immunofluorescent staining techniques, it was possible to show that NTHi biofilms, visible as bacterial aggregates surrounded by NTHi specific biofilm matrix components, formed within the smoke exposed animals infected with the parent strain, whilst no biofilms were detected in animals infected with mutant strains (Figure 4A). Further adding to the microscopy, RT-qPCR analysis of biofilm related genes showed increased expression of these genes of interest in parent strain infected animals (Figure 4B). Finally, we know that these biofilms are surrounded by activated neutrophils and exposed to neutrophil extracellular traps while within the smoke exposed lung (Figure 5). These neutrophil extracellular traps contribute significantly to the development of a highly oxidatively stressful environment and could thus be responsible for the failure of the thiol mutant strains to establish and infection within the susceptible animal model. Collectively, the *in vivo* data coincides and reinforces that *in vitro* work presented previously. Disruption of the thiol redox homeostasis pathway results in significant impairments in the ability of NTHi to establish a successful infection.

Taken together, our *in vitro* and *in vivo* work highlights importance of glutathione and thiol metabolism in controlling redox imbalances for NTHi, particularly within the biofilm mode of growth. Targeting these processes for biofilm-directed antimicrobials is an especially intriguing possibility for future work.

## Acknowledgments

The authors acknowledge helpful conversations and bioinformatics assistance from Al Claiborne (Wake Forest School of Medicine) as well as significant input and helpful discussions with colleagues. This work was supported by NIH research grants R21 AI133445 and R21 AI144507 to W.E.S, and RO1HL102371 awarded to A.G. B.C.H. was a trainee in the UAB Predoctoral Training Program in Lung Biology (NIH T32 HL134640).

